## Supplementary Figs. 1-8 and Table S1 for "Direct Regulation of the Voltage Sensor of HCN Channels by Membrane Lipid Compartmentalization"

This file includes:
Supplementary Figures 1-8 and the figure legends
Supplementary Table 1

Figure S1

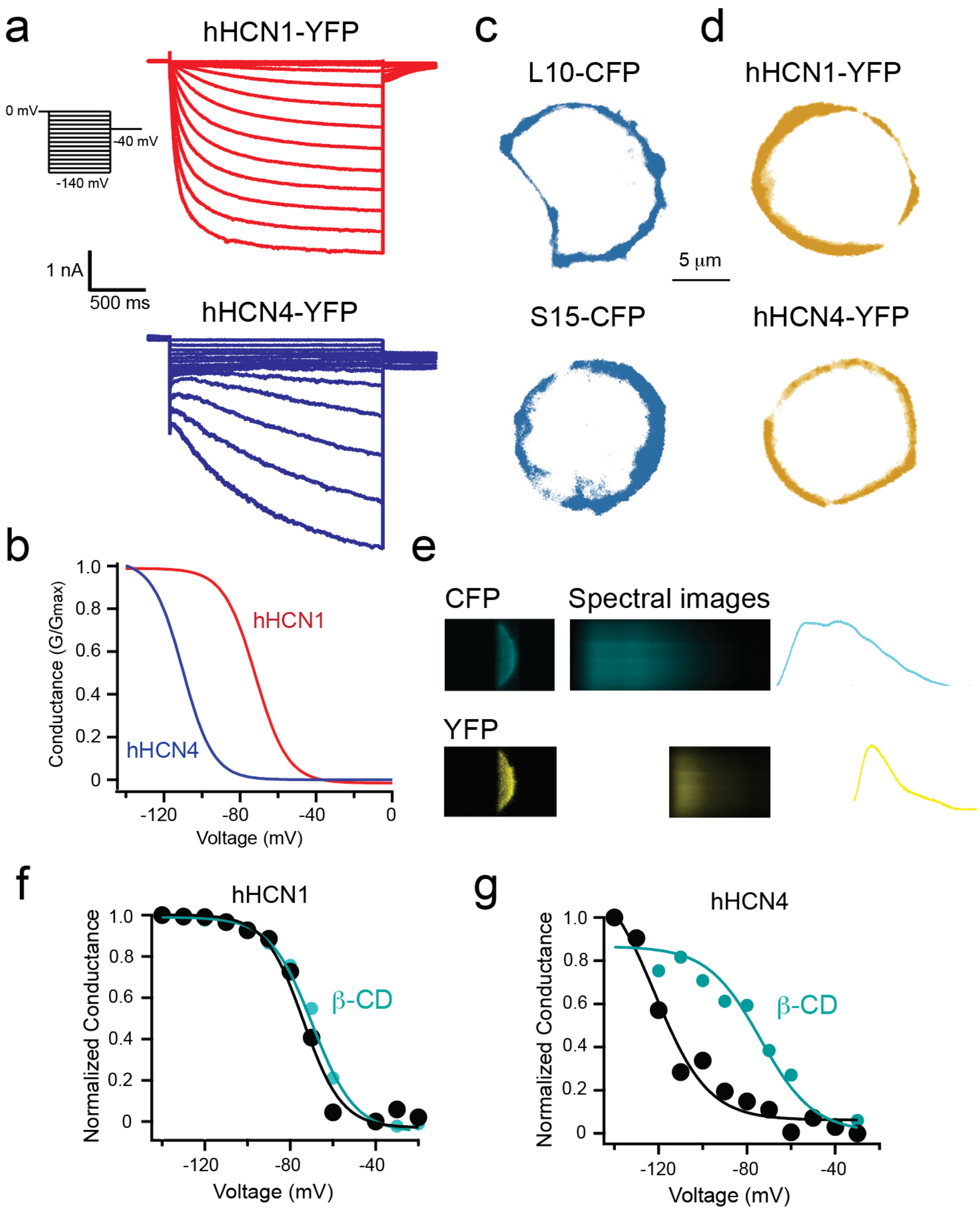


**Supplementary Figure 1. Properties of the constructs used for the spectral FRET between human HCN1-YFP, HCN4-YFP and L10-CFP, S15-CFP**

**a** Whole-cell ionic currents were recorded from hHCN1-YFP and hHCN4-YFP channels, elicited by a series of hyperpolarizing voltages ranging from 0 mV to -140 mV. **b** Conductance-voltage (G-V) relationships of HCN channels were estimated based on the tail current amplitude at the -40 mV voltage in the panel a. **c** and **d** Membrane-localized fluorescence was observed from

L10-CFP, S15-CFP, hHCN1-YFP, and hHCN4-YFP overexpressed in tsA201 cells. **e** Representative spectral images of CFP and YFP expressed cells were captured using a spectrograph equipped with a slit to select a narrow section of the epifluorescence from a single cell. Line scanning of the spectral images was performed to generate the emission spectra of CFP and YFP. Pseudo-colors were used for the images. **f** Effect of acute application of β-CD to disrupt ordered lipid domains on the G-V relationship of hHCN1-YFP construct, producing a minimal shift. **g** Effect of acute application of β-CD on the G-V relationship of hHCN1-YFP construct, producing a substantial depolarizing shift.

Figure S2

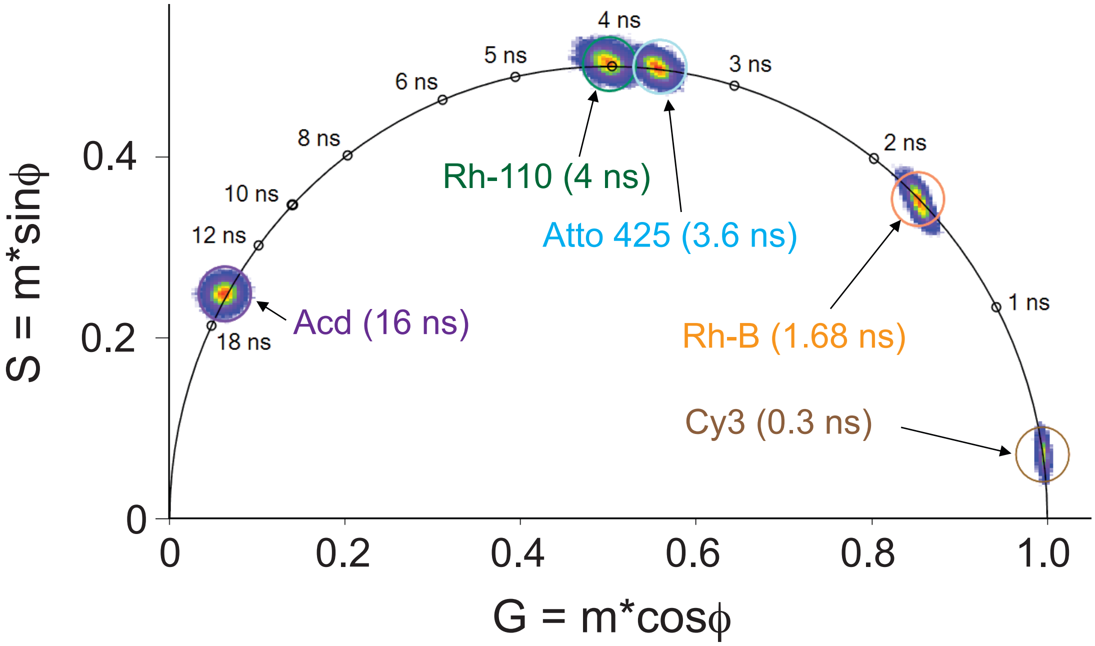


**Supplementary Figure 2. Phasor FLIM calibration using various fluorophores.**

Mono-exponential dyes, such as cyanine 3 (Cy3), rhodamine B (Rh-B), Atto 425, rhodamine 110 (Rh-110), and acridonylalanine (Acd), were selected for calibration purposes. These dyes exhibit lifetimes that align with the universal semi-circle on the phasor plot. Longer lifetimes are represented by an increase in phase angles (ϕ). The modulation (M) determines the distance of the phasor point from the origin (0, 0) in Cartesian coordinates. The calibration also corrects for the instrumental response function.

### Figure S3

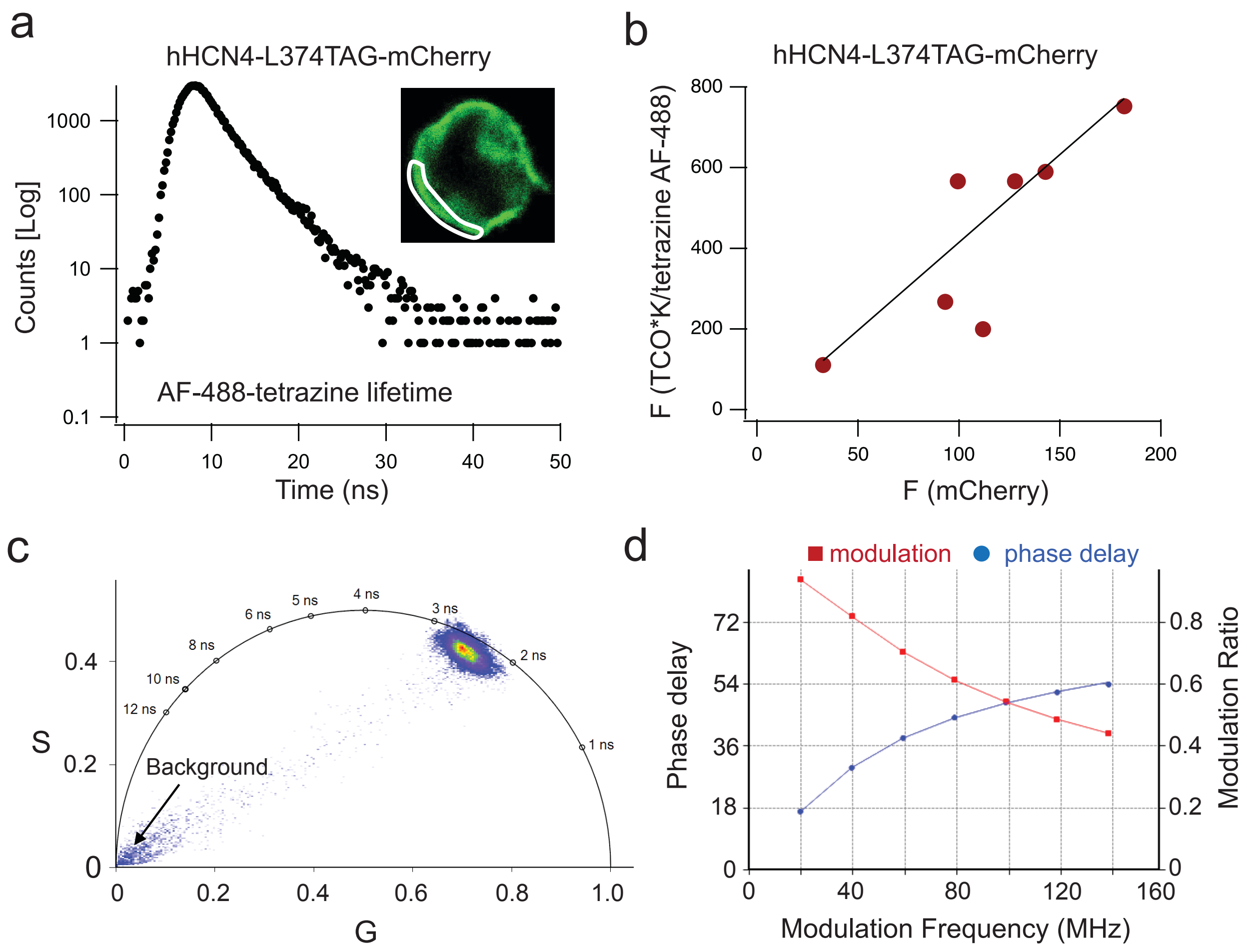


**Supplementary Figure 3. Properties of AF-488-tetrazine labeled to the TCO*K incorporated into the HCN voltage sensor.**

**a** Time-domain TCSPC measurement of the decay of photon emission of AF-488-tetrazine attached to the hHCN4-L374TCO*K-mCherry channels. The region under the cursor, localized to the membrane, was selected. **b** Correlation between the AF-488 fluorescence and the mCherry fluorescence from individual cells. **c** Phasor plot of the AF-488 from a single cell with the background lifetime species shown. No intensity-based filter was applied for this display. **d** Frequency-domain fitting of the AF-488-tetrazine from the same cell region as in the panel a. The modulation ratio decreases with the increase of the modulation frequency of the excitation laser (488 nm) while the phase delay increases with the modulation frequency.

Figure S4

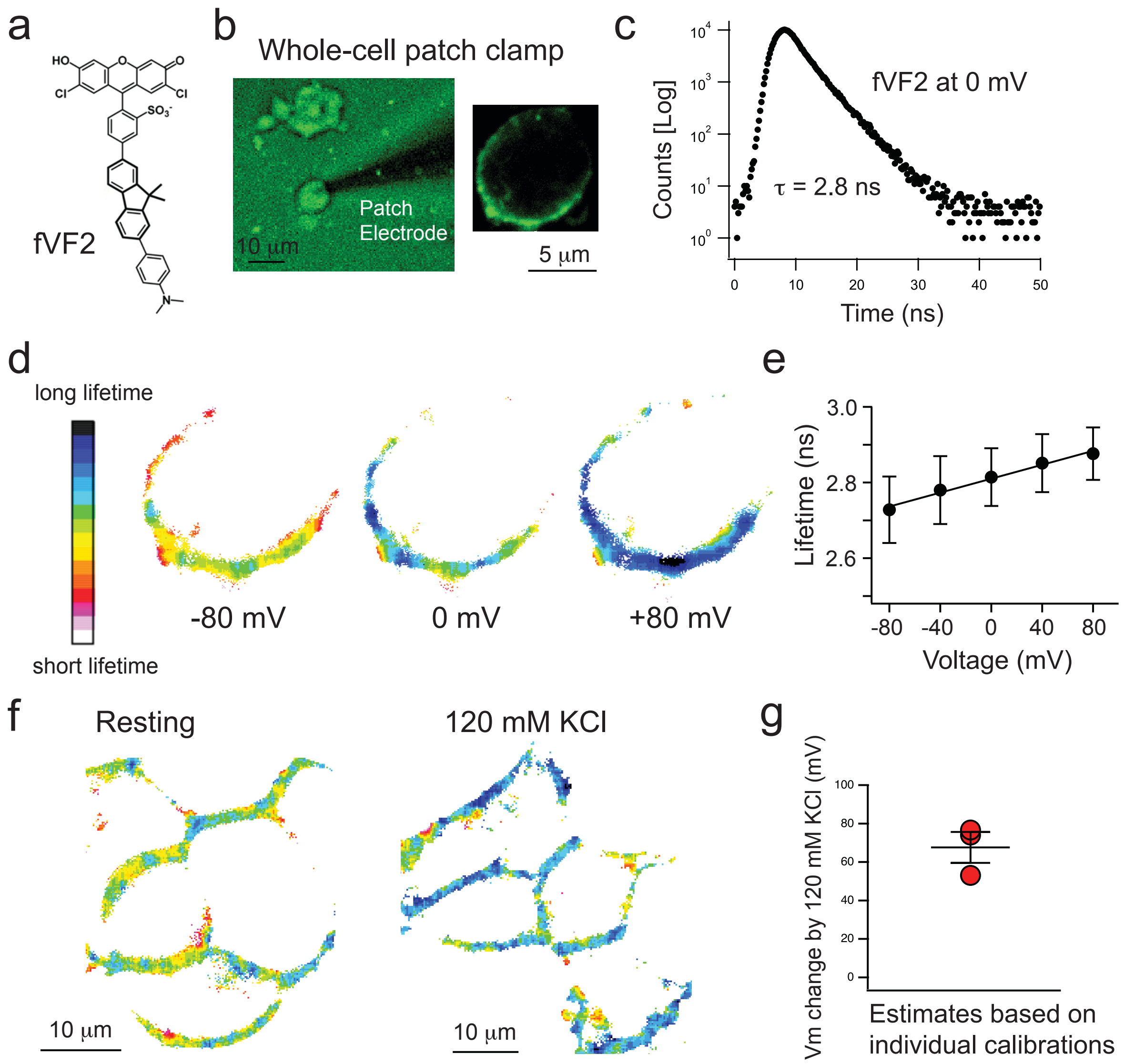


**Supplementary Figure 4. Estimating the degree of depolarization by 120 mM KCl using a voltage-sensitive dye reporting absolute membrane voltage.**

**a** Chemical structure of fVF2 dye, designed based on a photoinduced electron transfer mechanism, enables the estimation of absolute voltage through fluorescence lifetime measurement. **b** Whole-cell patch clamp recordings combined with FLIM were conducted on a tsA cell to measure the lifetimes of the fVF2 dye at different voltages. Bright-field illumination helped to display the shadow of the patch glass electrode. The image on the right shows an intensity-based representation of the same cell without bright-field illumination. **c** Time domain exponential decay of fVF2 emission of photons for the same cell in panel b, exhibiting a fraction-weighted average lifetime $\tau$ of 2.8 ns at 0 mV. Each pixel is analyzed using the same method of time domain lifetime fitting, generating a lifetime-based heatmap image as shown in the panel d.

**d** Heatmap images based on fluorescence lifetime measurements of the same cell from panel b are shown at various voltages. **e** A linear correlation between the membrane voltage and the lifetime of the fVF2 dye was observed, which is used for the calibration, n = 3. **f** Exemplar heatmap images based on fluorescence lifetime measurements of tsA cells are displayed, showing a comparison between the resting condition and after the application of 120 mM KCl. **g** Estimated changes in membrane voltage ($V_m$) following the application of 120 mM KCl are determined based on individual calibrations summarized in panel e.

Figure S5

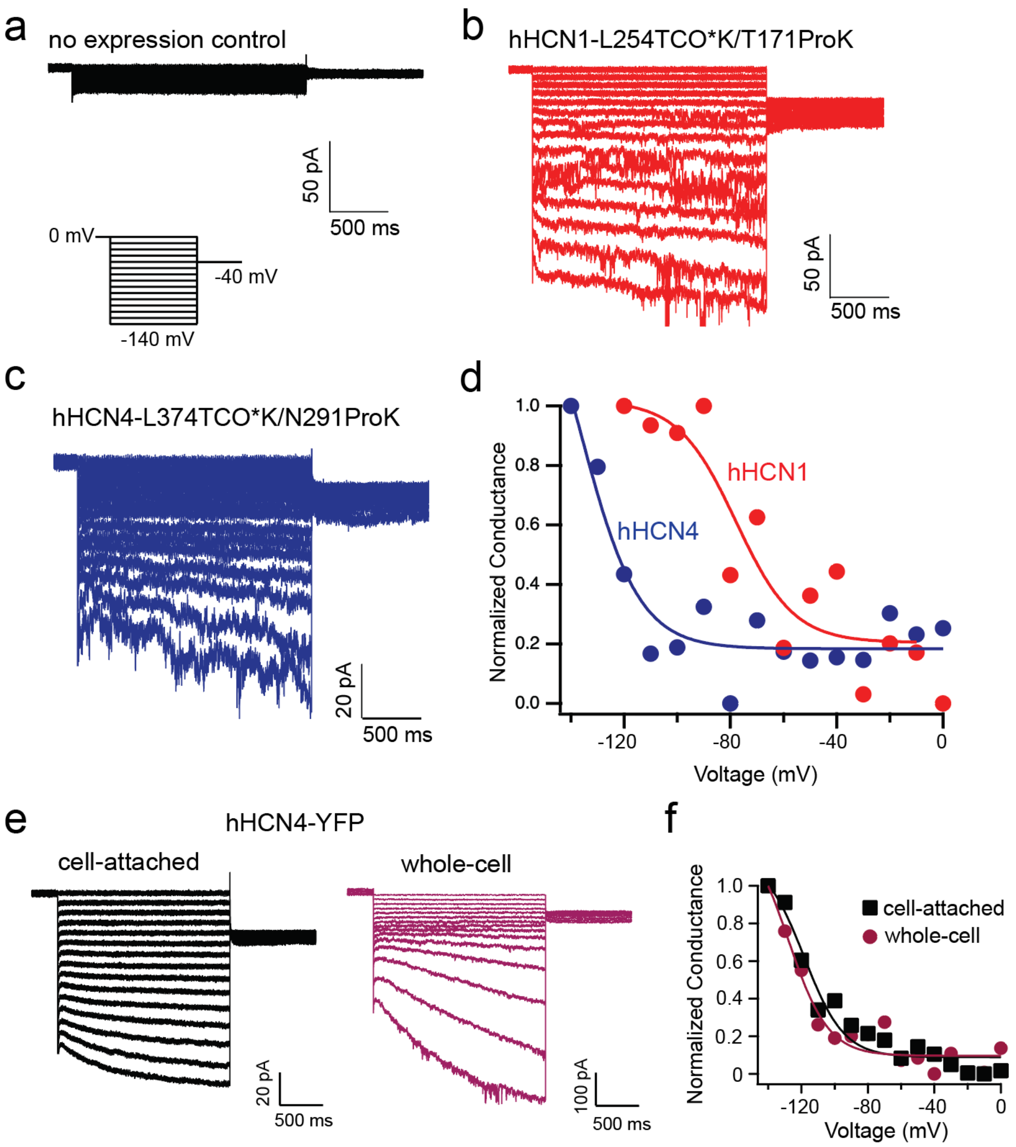


**Supplementary Figure 5. Properties of the hHCN channels with ncAAs incorporated using the dual codon-suppression strategy.**

**a, b** and **c** Ionic currents of hHCN1-L254TCO*K/T171ProK, hHCN4-L374TCO*K/N291Prok channels compared to the control with no channels expressed in tsA cells, recorded using the cell-attached patch clamp configuration. **d** Conductance-voltage (G-V) relationships of the HCN1 and HCN4 channels with ncAAs incorporated as shown in the panels b and c. **e** Ionic currents of hHCN4-YFP, recorded from the same tsA cell using the same voltage protocol as in the panel a, in the cell-attached or the whole-cell patch-clamp configuration. **f** Comparing the voltage dependence of hHCN4 channels measured in the panel e using the cell-attached and the whole-cell patch-clamp configurations, highlighting a similar G-V relationship.

#### Figure S6

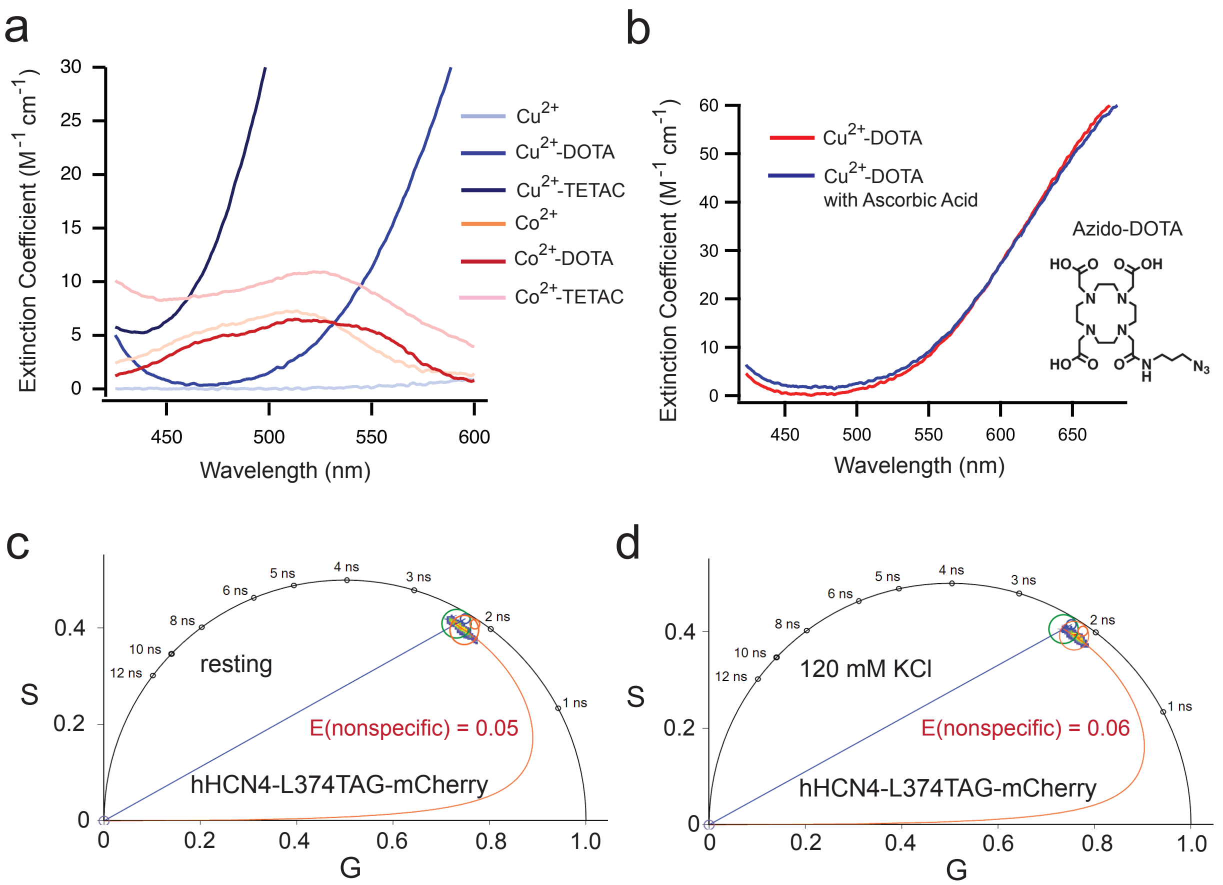


**Supplementary Figure 6. Properties of the molar absorption of transition metals**

**a** Absorption spectra of copper ($Cu^{2+}$) and cobalt ($Co^{2+}$) either in their free form or being chelated by DOTA or TETAC (1-(2-pyridin-2-yldisulfanyl)ethyl)-1,4,7,10-tetraazacyclododecane). **b** $Cu^{2+}$-DOTA molar absorption before or after mixing with an equal concentration (50 mM) of ascorbic acid for a half hour. **c** Representative phasor plot showing the nonspecific quenching of AF-488 caused by $Cu^{2+}$-DOTA using the hHCN4-L374TAG-mCherry construct, at the resting membrane condition. **d** Nonspecific quenching caused by $Cu^{2+}$-DOTA using the hHCN4-L374TAG-mCherry construct at the 120 mM KCl condition.

### Figure S7

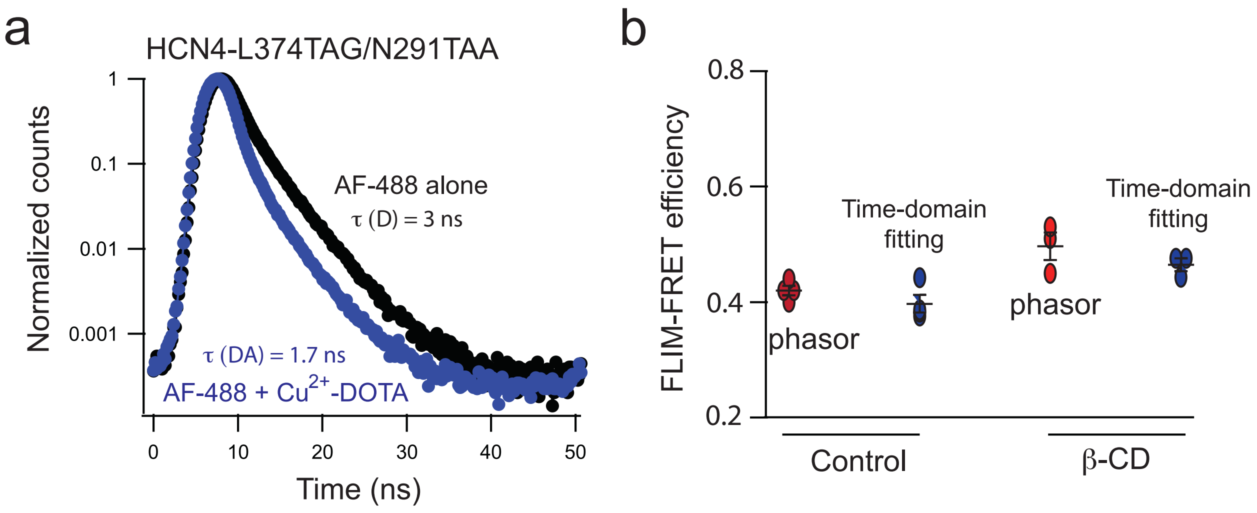


**Supplementary Figure 7. Comparing FRET efficiencies calculated using the FRET trajectory of the phasor plot versus the exponential fitting analysis.**

**a** Representative fluorescence decays of hHCN4-L374TCO*K/N291ProK expressed channels with or without applying $Cu^{2+}$-DOTA. The fluorescence lifetimes of the AF-488 with and without applying $Cu^{2+}$-DOTA obtained from the fitting analysis were 1.7 ns and 3 ns, respectively. **b** Comparing the calculated FRET efficiencies using the FRET trajectory function of the phasor plot versus using the double-exponential fitting.

Figure S8

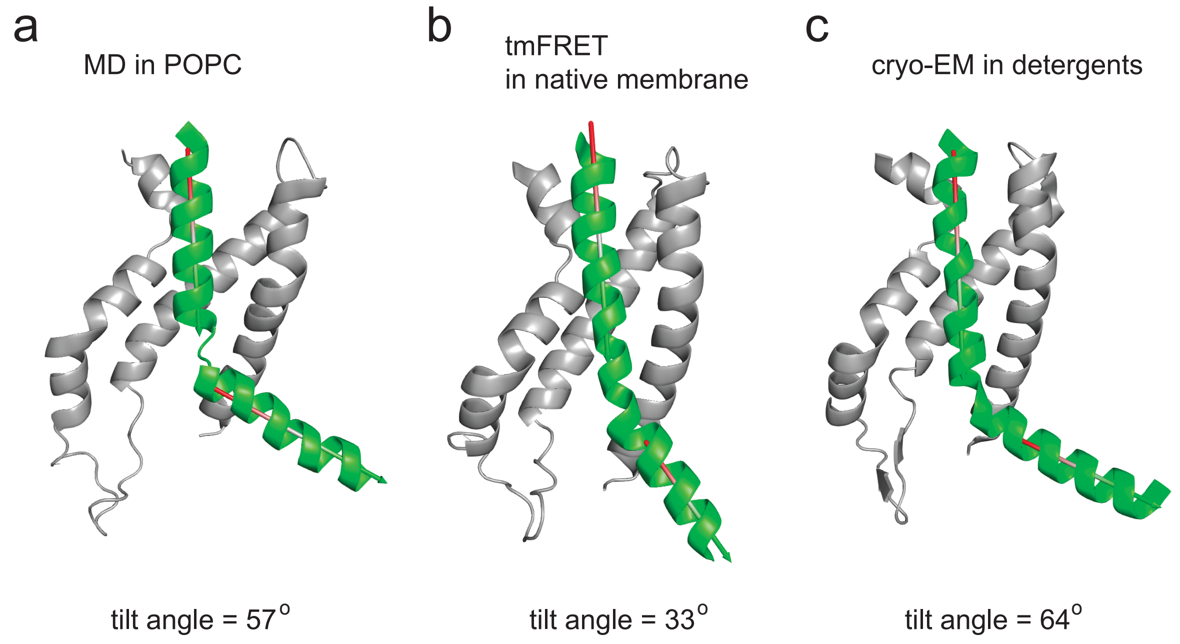


**Supplementary Figure 8. Varying degrees of tilting angles observed within the structural models of the S4 helix in HCN-VSD.**

Related to Figure 6: In **b**, a RosettaCM model that has been constrained by incorporating tmFRET-measured distances within the native Xenopus oocyte membrane. A hHCN4 homology model (SWISS-MODEL) was further built based on this RosettaCM model. Notably, this model exhibits a lesser degree of angle compared to **a**, which corresponds to a model generated through molecular dynamics (MD) simulation within a homogenous model membrane. Furthermore, **c** represents an alternative model, relying on a cryo-electron microscopy (cryo-EM) structure (PDB: 6UQF) of hHCN1 channel, solved in detergents. This model employs heavy metal-mediated cross-linking to stabilize the S4 helix in the downward conformation. The helix angels were estimated using the PyMOL plugin anglebetweenhelices.

Table S1: Estimated FRET efficiencies and distances between sites in the VSD of HCN1 and HCN4 channels using the Förster Convolved with Gaussian (FCG) relation.

| Conditions | HCN1-L254TAG, T171TAA pair (*$E_{FRET, corrected}$; Distance in* Å) | HCN1-I262TAG, T171TAA pair | HCN4 L374TAG, N291TAA pair | HCN4 I382TAG, N291TAA pair |
|---|---|---|---|---|
| Resting | 0.44; 17.2 | 0.38; 19 | 0.39; 18.7 | 0.36; 19.5 |
| Resting + β-CD | 0.39; 18.7 | 0.36; 19.6 | 0.48; 16.2 | 0.40; 18.4 |
| 120 mM KCl | 0.35; 19.8 | 0.47; 16.5 | 0.34; 20.1 | 0.46; 16.7 |
| 120 mM KCl + β-CD | 0.335; 20.3 | 0.46; 16.8 | 0.32; 20.8 | 0.44; 17.2 |
